# Optimizing On-the-Fly Probability Enhanced Sampling for Complex RNA Systems: Sampling Free Energy Surfaces of an H-Type Pseudoknot

**DOI:** 10.1101/2024.10.25.620366

**Authors:** Karim Malekzadeh, Gül H. Zerze

## Abstract

All-atom molecular dynamics (MD) simulations offer crucial insights into biomolecular dynamics, but inherent time scale constraints often limit their effectiveness. Advanced sampling techniques help overcome these limitations, enabling predictions of deeply rugged folding free energy surfaces (FES) of RNAs at atomistic resolution. The Multithermal-Multiumbrella On-the-Fly Probability Enhanced Sampling (MM-OPES) method, which combines temperature and collective variables (CVs) to accelerate sampling, has shown promise and cost-effectiveness. However, the applications have so far been limited to simpler RNA systems, such as stem-loops. In this study, we optimized the MM-OPES method to explore the FES of an H-type RNA pseudoknot, a more complex fundamental RNA folding unit. Through systematic exploration of CV combinations and temperature ranges, we identified an optimal strategy for both sampling and analysis. Our findings demonstrate that treating the native-like contacts in two stems as independent CVs and using a temperature range of 300–480 K provides the most effective sampling, while projections onto native Watson-Crick-type hydrogen bond CVs yield the best resolution FES prediction. Additionally, our sampling scheme also revealed various folding/unfolding pathways. This study provides practical insights and detailed decision-making strategies for adopting the MM-OPES method, facilitating its application to complex RNA structures at atomistic resolution.

**TOC Graphic:** 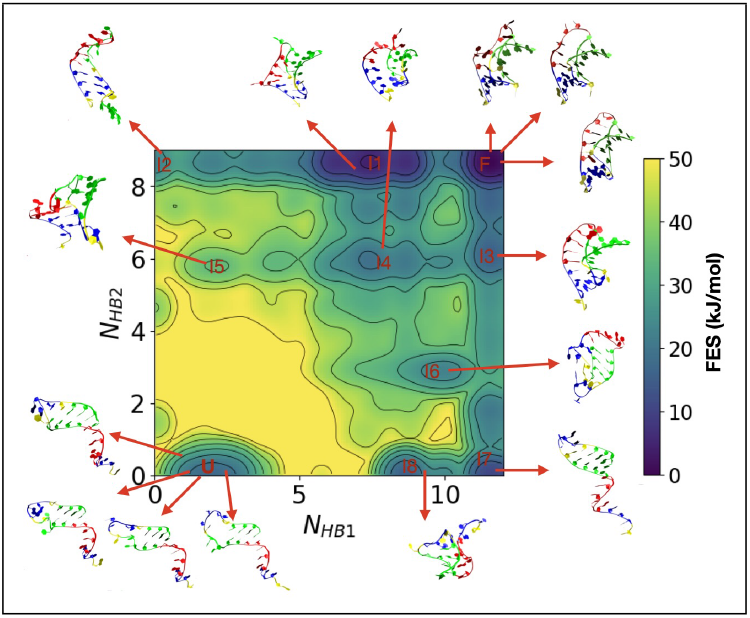

## Introduction

Atomistic molecular dynamics (MD) simulations are a staple for biomolecular studies, offering detailed insights into RNA conformational dynamics and stability in exquisite detail. However, these simulations face significant challenges in the presence of high energy barriers separating metastable states, which often is the case in RNA folding free energy surfaces (FES), resulting in rare transition events.^1,2^ Although sophisticated chip design has extended the limits of atomistic MD simulations, allowing significant increases in achievable simulation time,^3^ classical atomistic simulations still struggle to capture these transitions within accessible timescales. Advanced sampling techniques are essential to overcome these limitations, accelerating the exploration of phase space and providing information about relevant stable and metastable states. ^4,5^

Advanced sampling techniques in molecular simulations can be considered in two primary categories: tempering-based enhanced sampling and collective-variable-based enhanced sampling. In tempering-based methods, we leverage Boltzmann’s expression 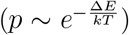 by increasing the system’s temperature to enhance sampling in high-energy regions. ^6–8^ Similarly, Eyring’s formula(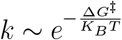, where *k* is the transition rate) suggests that simulations at elevated temperatures facilitate faster escape from local energy minima. The second category encompasses collective-variable-based sampling techniques. These methods are particularly effective for systems characterized by significant free energy barriers. They rely on an appropriate selection of relevant collective variables (CVs), which are functions of molecular coordinates. An appropriate CV (or set of CVs) for such sampling techniques typically captures important structural features or the slowest degrees of freedom of the system. We interchangeably referred to CVs as order parameters. In a subgroup of these CV-based techniques, bias potentials are applied to these chosen variables, guiding the simulations toward regions of interest in the conformational space.^9–15^ Successful implementation of CV-based techniques hinges on carefully selecting appropriate CVs that effectively capture the relevant aspects of the system’s behavior.

Pseudoknots are among the simplest yet most biologically significant RNA secondary structures that go beyond simple hairpins. They consist of interleaved base-paired stems and loops, which form a complex topology involving long-range interactions between different regions of the RNA sequence (e.g., Figure 1). This complexity results in more rugged FES than RNA hairpins, with a larger number of metastable intermediates that pose challenges for efficient sampling. Yet their structural intricacies are essential to their biological relevance, including their critical roles in trans-translation,^16^ ribosomal frameshifting,^17,18^ and viral replication.^19^ Capturing the folding pathways and energy landscapes of pseudoknots requires more sophisticated sampling methods to explore their conformational space fully.

**Figure 1.**
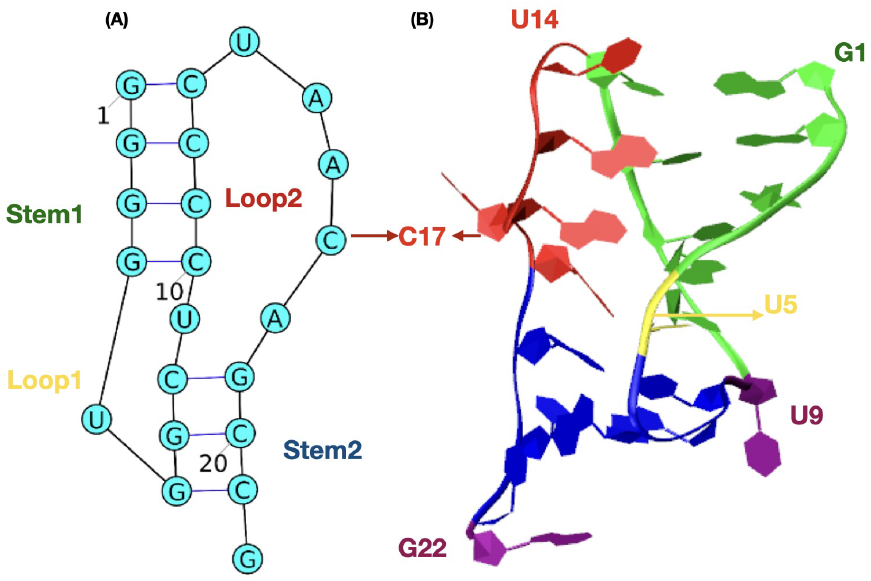
*Aquifex aeolicus* PK1 structure. (A) 2D structure of PK1. (B) 3D representation of the first frame of NMR structure (PDB ID: 2G1W) that shows interactions in stems and between loops and stems. Green, blue, red, and yellow colors represent stem 1, stem 2, loop 2, and loop 1, respectively. U9 and G22 nucleotides (which do not belong to any of the stems or loops) are represented with purple.

Recently, numerous investigations have been conducted to explore the configurational space of various RNA pseudoknots at the atomistic level using various methods aimed at accelerating sampling. Nguyen et al. conducted 8200 short independent MD simulations of various initial conditions (ICs) to study the folding, unfolding, misfolding, and conformational plasticity of the PLRV RNA pseudoknot.^20^ However, exploring all the relevant parts of the conformational landscape might be challenging with this technique due to the nontrivial nature of generating an unbiased set of ICs. Wang and co-authors used multiple conventional MD simulations to study the unfolding of the RNA pseudoknot within gene32 mRNA of bacteriophage T2 for the first time in atomistic detail. ^21^ However, since ensuring adequate sampling of the relevant parts of the phase space remains a challenge in this approach, more recently they used bias-exchange metadynamics (BEMD) to explore the folding FES of this RNA at atomistic detail in explicit solvent.^22^ BEMD involves concurrent biasing of predefined CV space in parallel running replicas but its order parameter-only nature might make BEMD simulations more susceptible to kinetic traps and hysteresis. Lazzeri et al. utilized ratchet-and-pawl molecular dynamics (rMD) simulations, a method that introduces a soft history-dependent biasing force to enhance the generation of productive folding trajectories towards a specified target structure. This approach was employed to generate the folding pathways of several RNA structures in the explicit solvent with low computational cost.^23^ However, a significant drawback of rMD simulations is that the generated trajectories tend to explore only a part of the energy surface. Specifically, they focus on the regions most likely involved in the productive pathways toward the target structure.^2^

Zerze et al. employed a hybrid approach combining parallel tempering and metadynamics to effectively sample all relevant parts of the folding FES of RNA tetraloops at the atomistic scale,^24^ which has proved useful to explore multiple free energy basins with similar stabilities. Subsequently, they adapted a multi-thermal, multi-umbrella variant of on-the-fly probability enhanced sampling (MM-OPES) approach, successfully reproducing the FES obtained from the combined parallel tempering and metadynamics simulations.^25^ MM-OPES proved to be nearly four times less costly and more straightforward to apply than its predecessor^25^ while providing the same benefits of combined temperature-based and order parameter-based sampling. However, despite its efficiency, applying and optimizing MM-OPES to characterize the FES of more complex RNA molecules, such as pseudoknots, where prior knowledge could be relatively limited, remained needed. This calls for further development and refinement of MM-OPES to tackle the structural characterization of FES of more complex RNA molecules with intricate underlying structures.

In this study, we aimed to adapt and optimize the MM-OPES method for an H-type RNA pseudoknot to improve sampling efficiency and gain deeper insights into RNA folding dynamics. The central challenge we addressed was devising the most appropriate set of CVs that accurately differentiate between various states and accelerate transition between the states, as well as optimizing temperature conditions that facilitate efficient sampling while minimizing any artificial effects on the distribution of conformations. We chose the “native-like contacts (Q)” as our primary order parameter, drawing on prior research where Zerze et al. successfully utilized this parameter in investigations of FES of both DNA^24^ and RNA tetraloops.^26^ To tailor this order parameter and MM-OPES for RNA pseudoknots, we explored different variations of Q based on distinct sets of atomic interactions within RNA structures for sampling purposes, coupled with varying temperature ranges. We projected the resulting FES onto various sets of order parameters, including those we used and those we did not use for biasing. We found that while some order parameters were more efficient for sampling the free energy surface, others could distinguish a larger number of metastable basins (although not as efficient in sampling). Notably, when projected onto two dimensions defined by the number of native hydrogen bonds (*N*_*HB*_) in stem 1 and stem 2 separately, we observed clearer distinctions between basins on the FES although the same order parameters were not as successful in improving the sampling’s efficiency.

Finally, we also uncovered multiple potential folding/unfolding mechanisms for the H-type pseudoknot via so-called reactive trajectory analysis based on extended biased trajectories. Notable mechanisms included step-wise initiation from either the first or second stem and cooperative mechanisms involving simultaneous formation/deformation of both stems. This detailed exploration sheds light on the intricate dynamics underlying RNA pseudoknot folding, contributing to a more comprehensive understanding of these complex structures.

## Material and Methods

### RNA Pseudoknot

Pseudoknot1 (PK1) in tmRNA of *Aquifex aeolicus* (figure 1)^27^ was chosen for our simulation. PK1 (5’-GGGGUGGCUCCCCUAACAGCCG-3’) is characterized as one of the smallest pseudoknot structures (22 nucleotides), consisting of two stems (which we will refer to as stem 1 and stem 2) and two loops (which we will refer to as loop 1 and loop 2). Stem 1 is formed by four base pairs: G1:C13, G2:C12, G3:C11, and G4:C10. Stem 2 comprises three base pairs: G6:C21, G7:C20, and C8:G19. Loop 1 is a single nucleotide loop with U5, while loop 2 spans U14 to A18. Out of its 22 nucleotides, only U9 and G22 do not participate in either of the stems or the loops (Figure 1). This concise architecture contributes to the efficient function of *Aquifex aeolicus* tmRNA, rendering it an intriguing subject warranting further investigation.

### Modeling

PK1 is modeled with the nucleic acid force field developed recently by Shaw and co-workers^28^ combined with the TIP4PD water model^29^ using the library files provided by Kuhrova et al,^30^ which will be referred to as DESRES ff hereafter. A single copy of PK1 was solvated in a truncated octahedron box. Na^+^ ions were added to provide electroneutrality, and the salt concentration was adjusted to 50 mM (NaCl). Ions were modeled by CHARMM22 parameters.^31^ Simulation input files were generated using AmberTools^32^ and then converted to a format compatible with GROMACS.

### Sampling

On-the-fly Probability Enhanced Sampling (OPES) was employed to sample the PK1 folding FES. This technique involves the introduction of a bias potential to sample from a modified target probability distribution, as outlined by Invernizzi et al.^12^ The target probability distribution is strategically designed to eliminate bottlenecks in the canonical Boltzmann distribution. Although OPES primarily operates as a collective variable-based technique, it possesses the versatility to sample generalized ensembles, such as the multicanonical or multithermal-multibaric ensembles.^11^ This is achieved by utilizing the potential energy and/or volume as collective variables, as demonstrated by Piaggi et al.,^33^ along with meticulously designed target distributions.^11^ OPES seamlessly accommodates the integration of generalized ensembles with CV-based enhanced sampling by incorporating the potential energy and a system-specific collective variable.^11,33^

In this work, we used a combination of a generalized ensemble and OPES, multithermal-multiumbrella OPES (MM-OPES) as previously adapted by Rahimi et al.^25^ MM-OPES is a variant of OPES, aiming to replicate and enhance both the effects of parallel tempering in the well-tempered ensemble and the effects of well-tempered metadynamics biasing. The latter involves diverse variations of the order parameter, constituting a multiumbrella expanded target distribution.^11^ We chose to use ‘native-like contacts (Q)’ as the order parameter to bias in the multiumbrella component, drawing inspiration from prior work. Notably, Zerze et al. previously employed this as an order parameter in their folding FES investigations on DNA^24^ and RNA tetraloop.^26^ The order parameter *Q* is defined by the following expression:

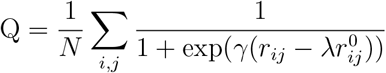

By adopting the generalized definition of *Q* as outlined in previous works,^34,35^ the summation in the equation encompasses *N* pairs, representing the total number of atomic pairs (*i, j*) considered in contact in the reference RNA state. When the *Q* is applied only for stems in the pseudoknot, any heavy atom (i.e., non-hydrogen) from the 3’ strand is considered in contact with a heavy atom from the 5’ strand if their interatomic distance is less than 5Å in the reference native structure. The same cutoff distance is used when *Q* is extended beyond the stems. The distances 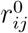 and *r*_*ij*_ denote the separations between atoms *i* and *j* in the reference native structure and the instantaneous configuration, respectively. In the smoothing function, *γ* is set to 50 nm^−1^, and the adjustable parameter *λ* is chosen as 1.5.^35^ Various adaptations of *Q* were employed as order parameters to enhance the sampling process, as discussed in the *Production* section.

To enhance the efficiency of our MM-OPES simulations, we implemented further parallelization through multiple walkers.^36^ In the multi-walker scheme, all walkers collectively sample the identical distribution, share a common bias potential, and collectively contribute to the free energy surface estimation.

### Equilibration

Following an energy minimization step, the system was equilibrated using the following strategy. First, an equilibration was conducted in the NVT ensemble for 100 ps with a time step of 2 fs using a velocity rescaling thermostat^37^ to keep the temperature at 300 K. Then, 100 ps equilibration in NPT ensemble was conducted with 2 fs time step, where Parrinello-Rahman barostat^38,39^ to maintain the pressure at 1 bar, and velocity rescaling thermostat^40^ for keeping the temperature at 300 K were used. This brief equilibration is followed by a 1 ns NPT equilibration for simulations at the running temperature and P = 1 bar prior to starting MM-OPES simulation. This step should not be skipped for appropriate biasing of potential energy. The temperature was maintained constant at any given run temperature using a velocity rescaling thermostat^37^ with a 1 ps time constant. Atmospheric pressure(1 bar) was maintained using an isotropic Parrinello-Rahman barostat^38,39^ with a time constant of 2 ps. Electrostatic interactions were calculated using the particle-mesh Ewald method^41^ with a real space cutoff distance of 1 nm. A cutoff distance of 1 nm was also used for the van der Waals interactions.

All simulations were performed using GROMACS (version 2021.4)^42–44^ patched with the PLUMED (version 2.8) enhanced sampling plugin.^45,46^

### Production

Since one of our goals in this manuscript is to identify the ideal set of CVs for sampling complex RNA folding FES using MM-OPES simulations, we set out to test different combinations of temperature ranges and CV sets (Table 1). The temperature range is divided into 5 groups based on the minimum and maximum temperature (*T*_*min*_-*T*_*Max*_). The CVs are organized into four sets: Set 1 involves biases applied separately to *Q*_*stem*1_ and *Q*_*stem*2_ (Table 2), treated as two distinct CVs. Set 2 involves biases applied separately to *Q*_*stem*1&*stem*2_ and *Q*_*tertiary*_ (Table 2) treated as two distinct CVs. Set 3 involves biasing added to *Q*_*all*_ (Table 2) as a single CV. Set 4 involves biasing added to only *Q*_*stem*1&*stem*2_ as a single CV.

**Table 1:** CV set and temperature combination tested in this work for exploring folding FES of PK1. These CV set and temperature range combinations yield a total of 12 independent simulations each with 8 walkers. The run temperatures (T_*run*_) are the arithmetic average of the minimum (T_*min*_) and maximum (T_*max*_) temperatures.

| CV set | Temperature ranges explored |
| --- | --- |
| Set 1: ($Q_{stem1}$ and $Q_{stem2}$) | 280-480, 290-530, 290-510, 290-490, 300-480 |
| Set 2: ($Q_{stem1\&stem2}$ and $Q_{tertiary}$) | 280-480, 290-530, 290-510, 290-490 |
| Set 3: ($Q_{all}$) | 280-480, 290-530 |
| Set 4: ($Q_{stem1\&stem2}$) | 280-480 |

**Table 2:** Collective variables (CV) used in this work for biasing and analysis.

| **Name** | **Definition** | **Purpose** |
| --- | --- | --- |
| $Q_{stem1}$ | # of heavy atom contacts in stem 1 of the native structure. | used for biasing (set 1) |
| $Q_{stem2}$ | # of heavy atom contacts in stem 2 of the native structure. | used for biasing (set 1) |
| $Q_{stem1\&stem2}$ | # of heavy atom contacts in stem 1 and stem 2 of the native structure. | used for biasing (set 2 and set 4) |
| $Q_{tertiary}$ | # of heavy atom contacts between loops and stems of the native structure (including U9 and G22). | used for biasing (set 2 ) |
| $Q_{all}$ | # of all heavy atom contacts of the native structure. | used for biasing (set 3) |
| $N_{HB1}$ | # of Watson-Crick type hydrogen bonds in stem 1 of the native structure | used for analysis (FES projection) |
| $N_{HB2}$ | # of Watson-Crick type hydrogen bonds in stem 2 of the native structure | used for analysis (FES projection) |
| $N_{HBs}$ | # of all Watson-Crick type hydrogen bonds of the native structure | used for analysis (FES projection) |
| $RMSD$ | All heavy atom root-mean-square distance from native structure | used for analysis (FES projection) |
| $eRMSD$ | Similarity between RNA structures and the native structure based on their three-dimensional structures | used for analysis (FES projection) |
| $R_g$ | All atoms Radius of gyration of structures | used for analysis (FES projection) |

### Analysis

We ran each production simulation for at least 1100 ns. After discarding 600 ns from the beginning of each simulation for equilibration, we calculated the unbiased FES projected on two dimensions (*CV* 1,*CV* 2) at temperature *T* ^*′*^ by using the formula,

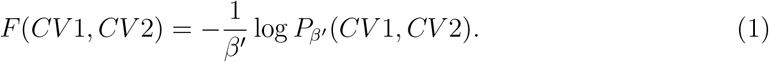

where *P*_*β*_*′* (*CV* 1, *CV* 2) is the unbiased probability of observing a microstate with a given value of *CV* 1 and *CV* 2 at a given inverse temperature *T* ^*′*^ = 1*/*(*k*_*B*_*β*^*′*^). In other words, *P*_*β*_*′* (*CV* 1, *CV* 2) is calculated as the weighted histogram of given *CV* 1 and *CV* 2 with the weights *w*(**R**) described as

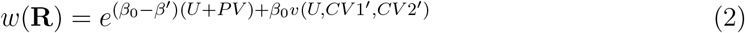

where *CV* 1^*′*^ and *CV* 2^*′*^ are the CVs that are used for biasing the given simulation (which may or may not be the same as CV1 and CV2), *v* is the bias potential,^11,25^ and *β*_0_ is the inverse running temperature *T*_0_ = 1*/*(*k*_*B*_*β*_0_).

*O*(**R**) is the ensemble average of any observable, and can be calculated at any inverse temperature, *β*^*′*^, of interest by using the partial weights of the microstates:

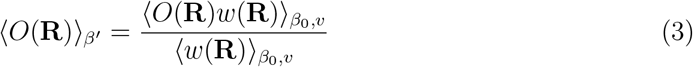

where ⟨·⟩_*β*_*′* is the average over the unbiased (isothermal-isobaric) ensemble at the temperature, *T* ^*′*^, of interest, *w*(**R**) is given by Eq. 2, and 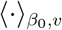 are the averages over the biased ensemble at the running temperature *T*_0_ with bias potential *v*.

We also performed a so-called reactive trajectory analysis which follows the biased trajectories in time and space to analyze the transitions between different states, following the approach developed by Zerze et al.^26^ We performed the reactive trajectory analysis on the FES projected on the number of hydrogen bonds present in stem 1 (denoted as *N*_*HB*1_) and those in stem 2 (denoted as *N*_*HB*2_) for each trajectory associated with individual walkers.

Structural clustering was performed based on structural similarity of the backbone heavy, i.e., non-hydrogen atoms, following the GROMOS algorithm^47^ using a 0.30 nm root-mean-square deviation (RMSD) cutoff distance. This cutoff is determined by systematically searching the ideal cutoff distance (Figure S1).

### Convergence

To investigate the convergence of the simulation and stability of the FES, we calculated the free energy differences between different states and the folded state (see XXX subsection for the definition of the states) using Eq. S1. Figure S2A shows the ✆*F* results as a function of simulation time. Notably, at approximately 1.5 microseconds, the free energy differences between different basins and the folded state stopped changing, signifying the convergence of our simulations.

To assess the statistical significance of the sample sizes at the 300 K for reweighting MM-OPES simulations, we computed the effective sample size (ESS) as a function of temperature using Eq. S2. The ESS can be interpreted as the number of samples that are relevant for reweighting at a given temperature. Figure S2B indicates the ESS is statistically significant enough (more than 2%)^48^ at lower temperatures, including 300 K (our reweighted temperature).

## Results and Discussions

### Efficiency of Conformational Sampling

To identify the ideal CV set and temperature range for exploring the FES of PK1, we first analyzed the time sampling of the *Q*_*stem*1&*stem*2_, one of the slow degrees of freedom for our system, for all CV sets and temperature ranges we tested (Figure 2). In metadynamics-based sampling, the success of the sampling (as well as the convergence) is typically assessed by the number of round trips between the upper and lower bounds of the slowest CV over the simulation course. An efficient metadynamics-based exploration yields many round trips in sufficiently long simulations, indicating a diffusive sampling between upper and lower boundaries. Atomistic simulations are typically slow due to the inherent high resolution and large number of degrees of freedom. Adding independent walkers (i.e., independent parallel running simulations), which contribute to the same free energy, partly addresses this concern. Since all walkers contribute to the same free energy surface in multiple walkers setting, ^36^ we plotted the time sampling of *Q*_*stem*1&*stem*2_ for all of the walkers on the same panel for a given simulation (Figure 2). The presence of extensive white space in the sampling between the upper and lower bounds of the CV typically results from (one or more) walkers being stuck in a metastable state for a long period and indicates lower diffusivity. We visually assessed the diffusivity of sampling in different panels of Figure 2 as discussed in the next paragraph.

**Figure 2.**
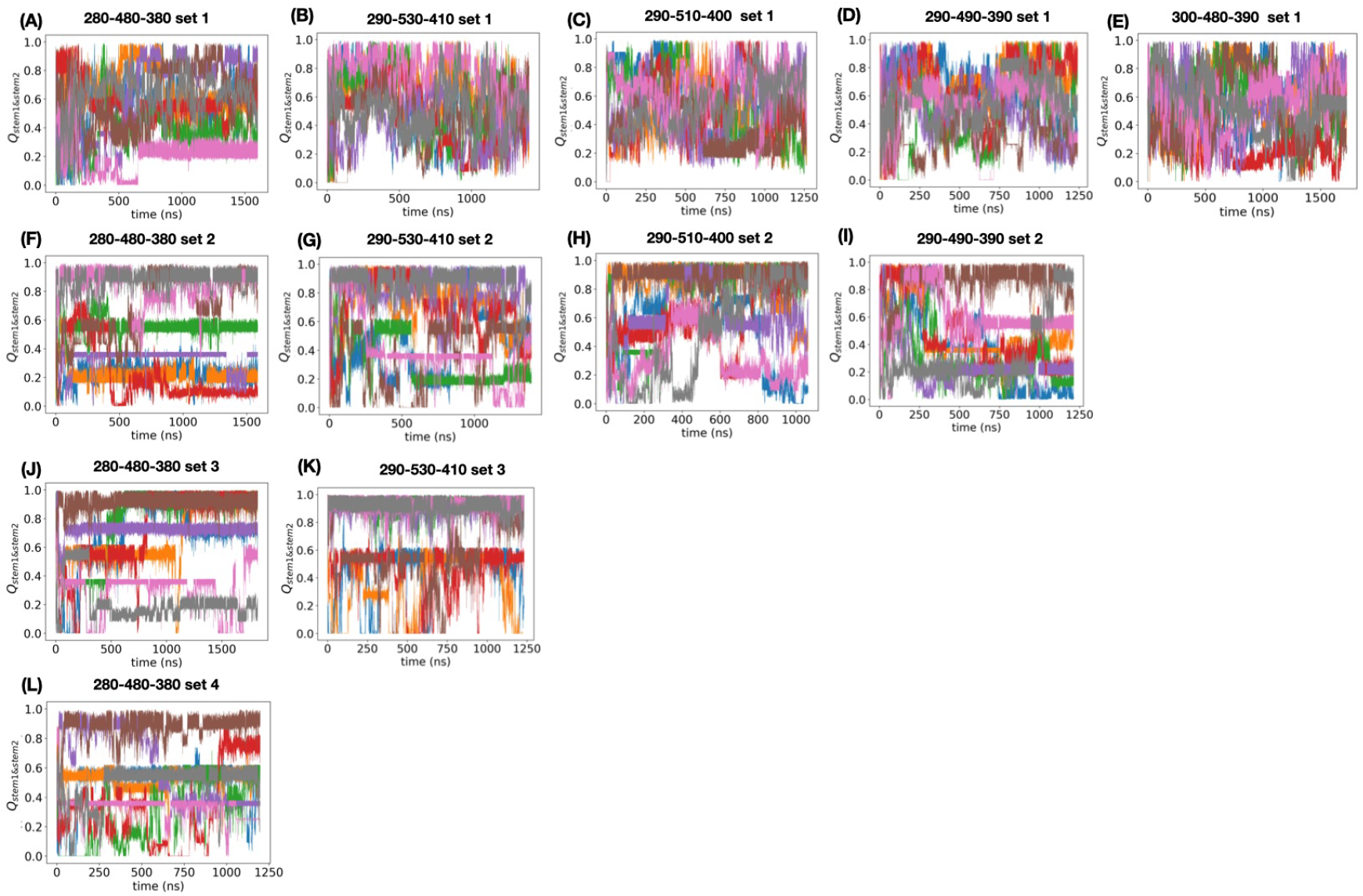
The efficiency of conformational sampling of our simulations tabulated in Table 1. under different conditions. Each panel depicts the sampling of *Q*_*stem*1&*stem*2_ as a function of time for all CV and temperature range combinations tested in this work (also see Table 1). The rows represent simulations that share the same CV set and columns represent simulations that share the same temperature range. Each plot is labeled with their temperature range, *T*_*min*_-*T*_*max*_-*T*_*run*_, and the CV set. Sampling of *Q*_*stem*1&*stem*2_ as a function time depicted for all of the walkers(each color represents a different walker) in each panel.

Temperature selections affect the diffusivity of sampling significantly (Figure 2). The multithermal ensemble explores the system’s potential energy between a minimum and a maximum temperature. These temperatures are indicated in the plot labels (with their corresponding run temperatures). Panels B, C, and D in Figure 2 show the effect of maximum temperature (when keeping the minimum temperature fixed) on the diffusivity of the sampling. Panel B in Figure 2 has the highest maximum temperature and has the most diffusive sampling. Panels A and E in Figure 2 compare different minimum temperatures. Elevated the minimum temperature (Panel E), yields increased diffusivity in sampling. In panels D and E, the running temperature (average of *T*_*min*_, and *T*_*max*_) remains fixed, but different temperature ranges are explored. Panel E has a narrower range of temperature (i.e., lower maximum and higher minimum temperature). Between panels D and E, E showed improved diffusivity, which we argue results from a larger minimum temperature. From this temperature exploration, we concluded that the higher the minimum and maximum temperatures are, the better the diffusivity is. However, we note that the temperature range also depends on the temperatures of interest for analysis. If the temperatures of interest fall outside the temperature range, the sampling is not likely to produce a statistically significant number of effective samples for the temperature of interest. Please see Rahimi et al.^25^ and Figure 4B for a relevant discussion. Since most biological phenomena and experiments occur at room temperature, we analyzed our simulations at 300 K and chose to limit our minimum temperature to 300 K for this reason.

Moving from the bottom to the top within each column in Figure 2 shows the comparison of different CV selections on sampling efficiency. Going from CV set 2 to set 1, we observe a notable improvement in the diffusivity of sampling. In CV set 1, we bias *Q*_*stem*1_ and *Q*_*stem*2_ as two separate CVs whereas in CV set 2, we have *Q*_*stem*1&*stem*2_, which combines *Q*_*stem*1_ and *Q*_*stem*2_ into a single CV, and we have *Q*_*tertiary*_, which contains long-range atomistic contacts between loops and stems. Further detailed definitions of these CVs are given in Table 2. The improved diffusivity in CV set 1, compared to CV set 2, emphasizes the importance of treating contacts in each stem of the pseudoknot as orthogonal reaction coordinates, instead of combining them into one. In rows 3 and 4 in Figure 2, CV set 3 and CV set 4, only one CV, *Q*_*all*_ (CV set 3) and *Q*_*stem*1&*stem*2_ (CV set 4), respectively, is considered for multiumbrella sampling. Diffusivity in these rows is lower than simulations in rows 1 and 2 when considering two CVs for accelerating sampling. This indicates that one CV is insufficient to accelerate the escape from metastable basins for efficient sampling of the conformational landscape. From this CV set comparison, we concluded that the set that treats each stem as an orthogonal reaction coordinate has the best efficiency in sampling the relevant conformational space of this pseudoknot.

Based on our analysis of various temperature ranges and CV sets, we selected the temperature range 300-480 K with CV set 1, *Q*_*stem*1_ and *Q*_*stem*2_, (the simulation whose efficiency of conformational sampling is presented in panel E in Figure 2) for the further in-depth analysis presented below.

### Free Energy Surfaces

Figure 3 illustrates the folding free energy surfaces (FESs) of PK1 projected onto different collective variables by reweighting the simulation from Figure 2E at 300K. Each CV has a different capability of distinguishing metastable states of this system. We characterized the folded state (F) based on its heavy atom RMSD (using reference structure PDB ID: 2G1W) *<*0.65 nm and the fraction of native-like contacts in stems *>* 0.8. The unfolded (U) state is labeled as the state that has the least number of Watson-Crick (WC) type hydrogen bonds. Intermediate (Ix) states are the metastable states between F and U with some WC-type hydrogen bonds in stems and native-like contacts. While the unfolded and intermediate states vary relatively largely between different projections, the folded basin is shared between all projections with a minimum of 80% population overlap between all projections.

**Figure 3.**
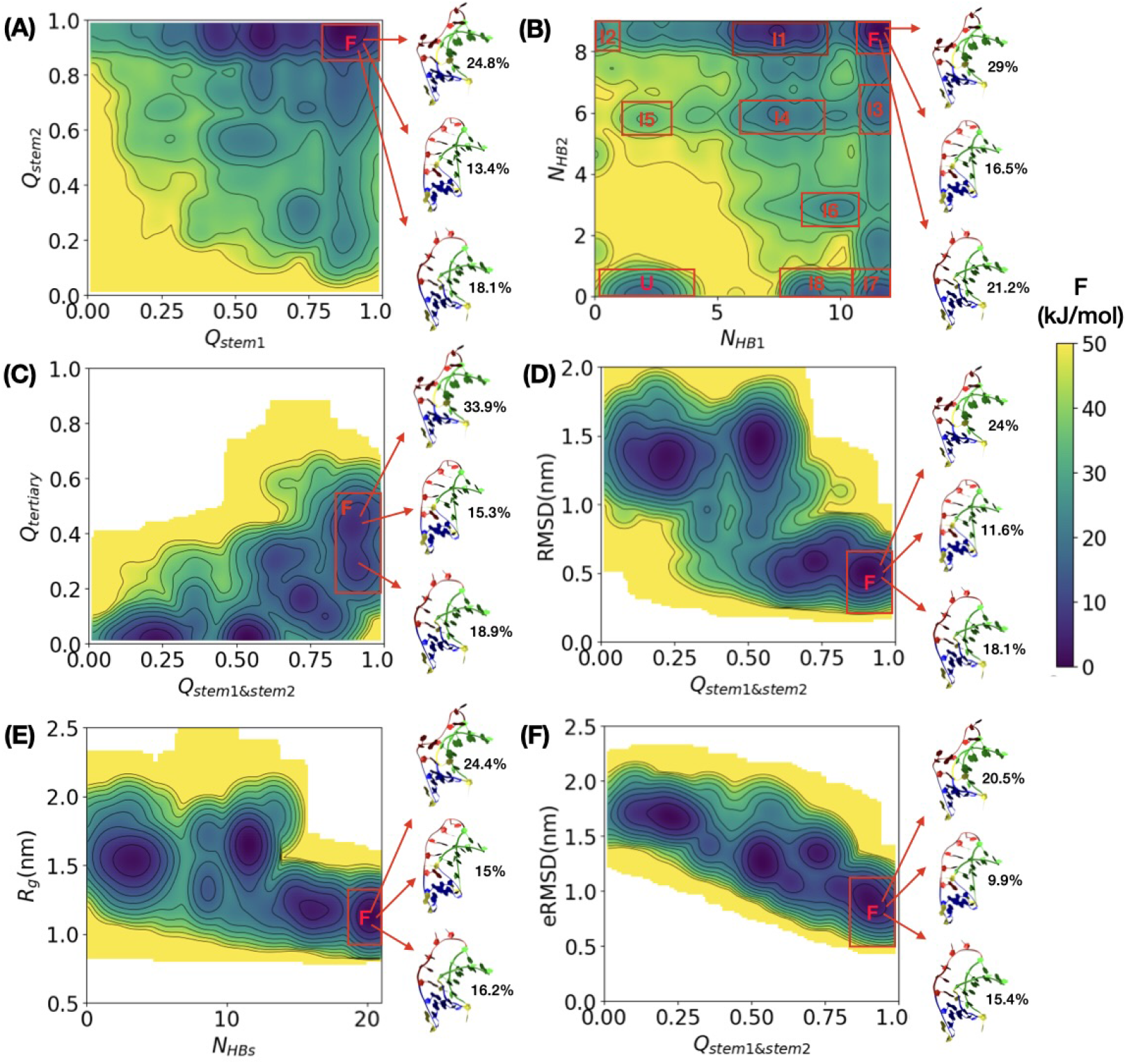
Folding free energy surfaces (FES) of PK1 is projected on multiple selected CV combinations at 300 K. Unbiased FES was calculated by reweighing MM-OPES simulations conducted at 300-480 K temperature range, employing CVs set 1. All FES are reweighted at 300 K. Panel A depicts the FES projected on the CVs used for accelerating sampling, all other panels show the FESs projected on different CVs (not used in the biasing). The folded state is labeled as F in all panels and the three most populated clusters of folded state are indicated with arrows with their percentage population. Unfolded, and intermediate states are labeled as U, and Ix, respectively, in panel B.

We, first, projected the FES onto the CVs used to accelerate the simulation (Figure 3A). However, compared to panel B, this CV combination fails to resolve all relevant metastable basins, yielding fewer, more degenerate intermediate basins although their F basins share 99% population between panel A and B projections.

Panel B displays the FES projected on the number of Watson-Crick-type hydrogen bonds in stem 1 (*N*_*HB*1_) and stem 2 (*N*_*HB*2_) are expected to form in the native structure (PDB ID: 2G1W). These CVs combination effectively separates the metastable basins on the FES and maximizes the number of metastable basins, offering a distinct and efficient representation compared to other alternatives presented in Figure 3. Accordingly, we labeled all intermediate states and the unfolded state on Panel B. While this hydrogen bond-based CV set is the most effective in maximizing the number of distinguishable metastable basins, it was not equally effective in accelerating the sampling as we have shown previously.^24^ To test this, we conducted further simulations where we biased *N*_*HB*1_ and *N*_*HB*2_ as two orthogonal CVs using the ideal temperature range we identified (300-480 K). This CV set was indeed insufficient to accelerate the simulation and enable the system to escape from the metastable states as evident from the time evolution of *Q*_*stem*1&*stem*2_, which shows some walkers being trapped at metastable intermediates (Figure S3).

Structural clustering analysis showed that the three most populating clusters (with slightly varied percent populations as demonstrated in Figure 3) are the same for the F basins of all panels. The primary distinction between these clusters lies in the backbone twisting of loop 2. This twist is quantified by backbone dihedral angles. We found the most prominent differences in the *α*_*C*17_ and *ζ*_*U*14_ angles whose distributions can be seen in Figure S4 for the folded state. This backbone twisting influences the contacts between residues in loop 2 and stem 1. Given that the tertiary interactions in PK1 are predominantly governed by the interactions between loop 2 and the minor groove of stem 1,^27^ these structural changes directly impact *Q*_*tertiary*_. Since *Q*_*tertiary*_ can capture these conformational differences in the folded state, FES is projected onto *Q*_*stem*1&*stem*2_ and *Q*_*tertiary*_ in panel C. This projection reveals two distinct (sub)basins within the folded state, separated by an energy barrier of less than 5 kJ/mol. The upper (sub)basin contains the first and third most populated structural cluster, while the lower (sub)basin is associated with the second most populated structural cluster. Notably, these (sub)basins are unique to panel C and do not appear in other panels. We hypothesize that the structural differences in the folded state, similar to what we observe here, (arising from the backbone twisting of loop 2 that modifies loop 2 and stem 1 tertiary contacts) could explain the experimentally observed functional changes (altered efficiency of the frameshifting) to a frameshifting pseudoknot^49^ as suggested in other structural studies as well.^18,20^

In panel E, the FES is projected onto all Watson-Crick type hydrogen bonds of the native structure (*N*_*HBs*_) and the radius of gyration (*R*_*g*_). This panel provides useful insight into compactness with respect to hydrogen bonding, but the lack of distinct energy wells especially across *R*_*g*_ makes it less ideal for free energy projections. Finally, panel F depicts the FES as a function of eRMSD^50,51^(calculated for nucleotides in stems) and *Q*_*stem*1&*stem*2_. This panel resembles Panel D but uses eRMSD, which is a metric developed for measuring distances between three-dimensional RNA structures. The energy landscape here shows a (negative) correlation, where high *Q*_*stem*1&*stem*2_ values consistently correlate with low eRMSD, indicating a strong coupling between the formation of native-like stem contacts and the proper assembly of the three-dimensional stem structure. However, compared to panel B, it cannot maximize the number of metastable basins suggesting that they may not be ideal order parameters for detailed analysis.

Our analysis demonstrates that while different collective variables (CVs) offer complementary benefits to the folding landscape of the PK1 pseudoknot, no single CV set is universally optimal. Although *Q*_*stem*1_ and *Q*_*stem*2_ are superior in reaching a diffusive sampling (Figure 2E), projections based on hydrogen bonds (Figure 3B), provide the best separation of metastable basins compared to all other CVs (including *Q*_*stems*_ shown in Figure 3A). Despite providing the best separation, we found that the hydrogen bond-based CVs are insufficient for enhancing sampling efficiency. Additionally, projections such as eRMSD and RMSD capture structural similarity but lack the ability to maximize basin resolution. Meanwhile, projections on *Q*_*tertiary*_ can distinguish potentially different functional forms of the F states but cannot identify most intermediate states. Ultimately, this analysis emphasizes the importance of carefully selecting order parameters to balance sampling efficiency with structural resolution, particularly when characterizing complex RNA structures like H-type pseudoknots.

### Analysis of Reactive Trajectories

The folding FES (Figure 3B) of PK1 reveals the presence of multiple metastable basins, indicating that the pseudoknot folding/unfolding could follow several distinct paths visiting different metastable intermediates. To comprehensively explore all possible folding/unfolding pathways in our simulations, we analyzed each trajectory by identifying all sub-trajectories that start from the unfolded state, U, and ultimately reach the folded state, F (Figure 4A), and vice versa (i.e., from F to U) (Figure 4B). This so-called “reactive trajectory” analysis aims to gain insights into the folding/unfolding pathways and determine how intermediate states are encountered along these pathways.

**Figure 4.**
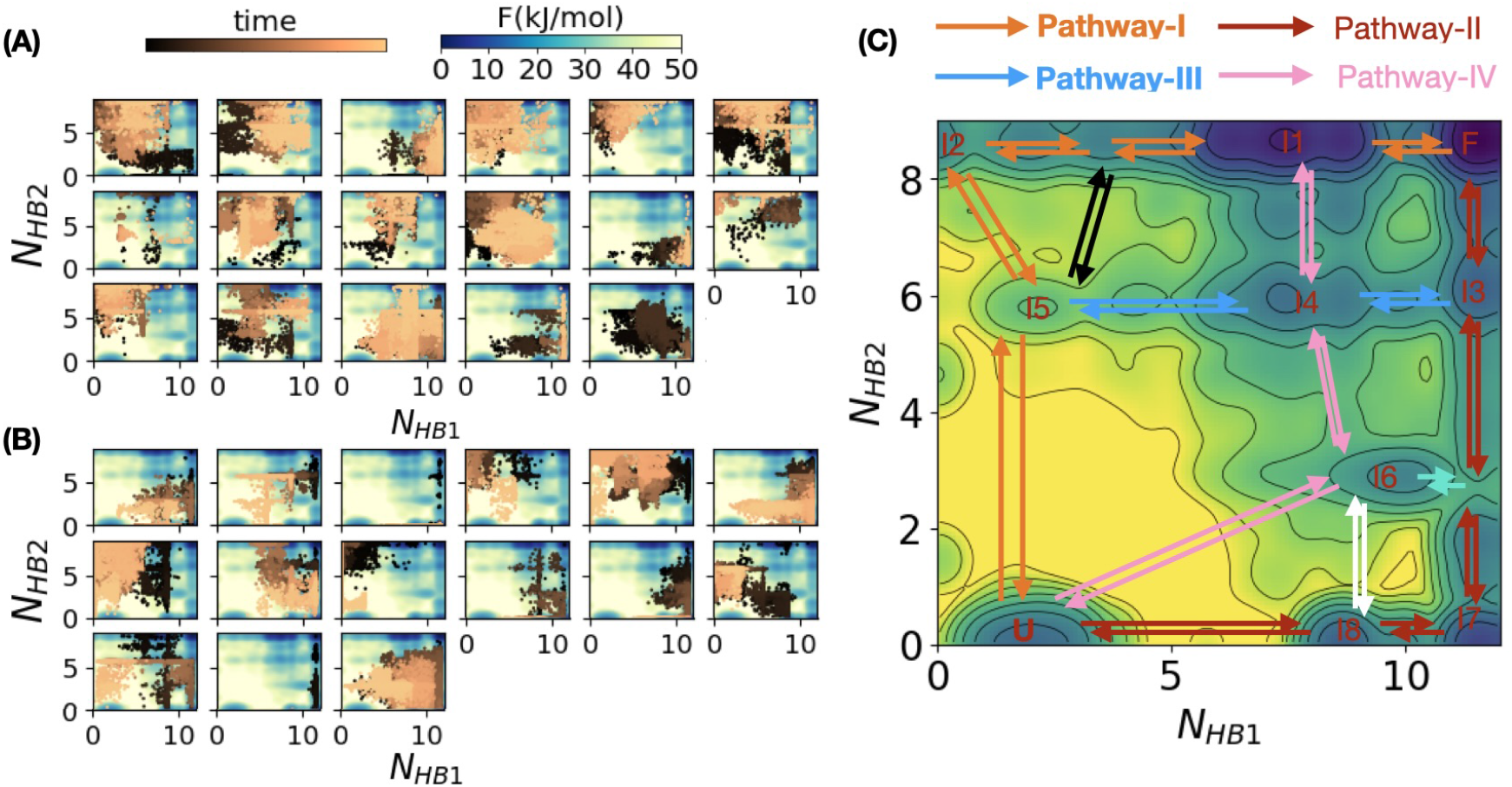
Path analysis of transitions between different states. Panel A shows the paths (i.e., the visited points as a function of time) on the same *N*_*HB*1_ vs *N*_*HB*2_ FES during U to F transitions (i.e., transitions starting from U and landing in F) for all U to F transitions in simulation conducted at 300-480 K temperature range, employing CVs set 1. A total of 17 U to F transitions are found. The color bar labeled “time” represents the progression along each transition, with dark brown indicating the beginning of a transition and yellow representing the end. Panel B shows the path on the same FES during F to U transitions (15 F to U transitions in total). Panel C reveals multiple potential folding and unfolding pathways, identified through the analysis of sub-trajectories in Panels A and B. Several key pathways are color-coded at the top of the panel. These pathways, either individually or through hybrid mechanisms, contribute to the overall folding and unfolding process.

The analysis of all these sub-trajectories revealed several folding /unfolding pathways (Figure 4C) for this RNA pseudoknot. Please note that in this context, the term “pathway” is used to refer to the thermodynamic routes in the FES. ^22^ This usage does not refer to the kinetic processes or transition rates along these routes.

Two primary pathways (Pathway-I and II) dominate the folding and unfolding of PK1. In Pathway-I, stem 2 forms before stem 1 during folding, and stem 1 deforms first during unfolding. Pathway-II follows the opposite sequence, where stem 1 forms before stem 2 during folding, and deforming last during unfolding. These step-wise parallel mechanisms found here align with the folding mechanism previously reported for the PK5 pseudoknot by Cao et al.^52^ and are consistent with the characteristics of the MMTV PK folding mechanism elucidated through combined laser temperature-jump experiments and coarse-grained simulations.^53^ These step-wise mechanisms also align with the findings of Lazzeri et al. who used ratchet-and-pawl MD (rMD) to obtain the folding mechanisms of the PK1 by starting from unfolded configurations.^23^

We also found alternative pathways where both stems of PK1 pseudoknot simultaneously fold and unfold, corresponding to pathway-III and pathway-IV (Figure 4C) that embody a cooperative mechanism where both stems form or deform together. Finally, we also found hybrid mechanisms, i.e., combined step-wise and cooperative mechanisms. For instance, folding might proceed starting from Pathway-IV from U to I6 then continuing to F from Pathway-II, and vice versa as shown by green arrows on Figure 4C. Similar hybrid mechanism jumps between step-wise and cooperative mechanisms could also be seen by black and white arrows in Figure 4C.

These multiple folding/unfolding mechanisms are consistent with the reported folding mechanisms of pseudoknot within gene 32 mRNA of bacteriophage T2, ^21,22^ and PLRV frameshifting pseudoknot.^20^ Our findings reveal striking similarities in the folding and unfolding pathways of this H-type pseudoknot. The agreement across independent studies underscores the robustness and reliability of the observed behaviors in the PK1 structure.

## Conclusion

In this work, we have adapted a novel advanced sampling technique, Multithermal-Multiumbrella On-the-Fly Probability Enhanced Sampling (MM-OPES), to study folding free energy landscapes and folding/unfolding pathways of an H-type RNA pseudoknot.

Through a systematic exploration of temperature ranges and CVs, we optimized the MM-OPES for H-type RNA pseudoknots to facilitate escape from energy traps and maximize the number of metastable states. Our extensive CV analysis revealed that no single CV set is universally optimal for studying the folding free energy surfaces of PK1. While *Q*_*stems*_ proved most effective in accelerating sampling, WC-type hydrogen bonds best separated metastable states and maximized the number of metastable basins.

The efficient sampling achieved by MM-OPES, with repeated transitions between folded and unfolded states, enabled us to trace thermodynamic paths along reactive trajectories. This analysis uncovered multiple folding and unfolding mechanisms, including step-wise, cooperative, and hybrid pathways, aligning with prior experimental observations.

MM-OPES shows significant potential for atomic-level characterization of complex equilibrium ensembles in biomolecular systems. However, successful application requires careful selection of CVs and optimization of temperature ranges for effective exploration. Our study provides a comprehensive guide for adapting MM-OPES to complex RNA folds, offering a valuable framework for future research on more intricate RNA structures of interest to the community.

## Acknowledgement

This work was supported by funding from the Cancer Prevention and Research Institute of Texas (CPRIT) award RR220008 and the Welch Foundation (Award E-2221 and Catalyst Center for Advanced Bioactive Materials Crystallization Award V-E-0001). The simulations presented in this work were performed on the computational resources provided by the Hewlett-Packard Enterprise Data Science Institute at the University of Houston.

## Supporting Information Available

A plot of sampling of *Q*_*stem*1&*stem*2_ as a function of time for simulation where *N*_*HB*1_ and *N*_*HB*2_ CVs are biased, the heatmap of the probability distribution of backbone conformation, statistical evaluation for convergence and effective sample size of MM-OPES, and analysis of choosing proper cutoff for clustering analysis are available.

## Supporting Text

### Statistical evaluation

In Figure 3B, multiple intermediate states as well as the folded and unfolded states were identified. To investigate the convergence of the simulation and stability of the FES, we calculated the free energy differences between different states and folded state (✆*F*_*F U*_ = *F*_*folded*_ − *F*_*unfolded*_, ✆*F*_*F Ix*_ = *F*_*folded*_ − *F*_*intermediate*_) (*x* being any of the intermediate states) follows:^1^

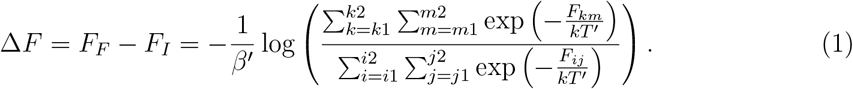

Indices *m*_1_&*m*_2_ and *k*_1_&*k*_2_ are the boundaries of the discretized *N*_*HB*1_ and *N*_*HB*2_ coordinates for the folded state; i.e., *F*_*km*_ is the free energy corresponding to a given bin on the *N*_*HB*1_ and *N*_*HB*2_ coordinates of the folded state. Indices *i*_1_&*i*_2_ and *j*_1_&*j*_2_ are the boundaries of the discretized *N*_*HB*1_ and *N*_*HB*2_ coordinate of any intermediate or unfolded states; i.e., *F*_*ij*_ is the free energy corresponding to a given bin on the *N*_*HB*1_ and *N*_*HB*2_ coordinates of any intermediate or unfolded states. The ✆*F* results are shown as a function of simulation time in Figure S2A.

We also calculated the effective sample size (ESS) at 300 K to assess the statistical significance of the reweighted FES at 300 K as follows:^2–4^

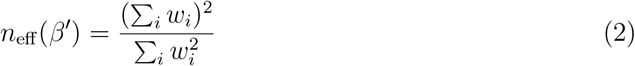

where 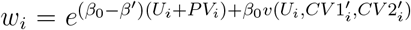 are the importance sampling weights of the *i*-th configuration of the trajectory, and 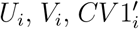, and 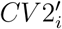 are the potential energy, volume, *Q*_*stem*1_ and *Q*_*stem*2_ for the same configuration. Figure S2B shows the ESS is statistically significant enough (larger than 2%)^5^ at lower temperatures for the MM-OPES simulation conducted at 300-480 K temperature range, employing CVs set 1.

## Supporting Figures

**Figure S1:**
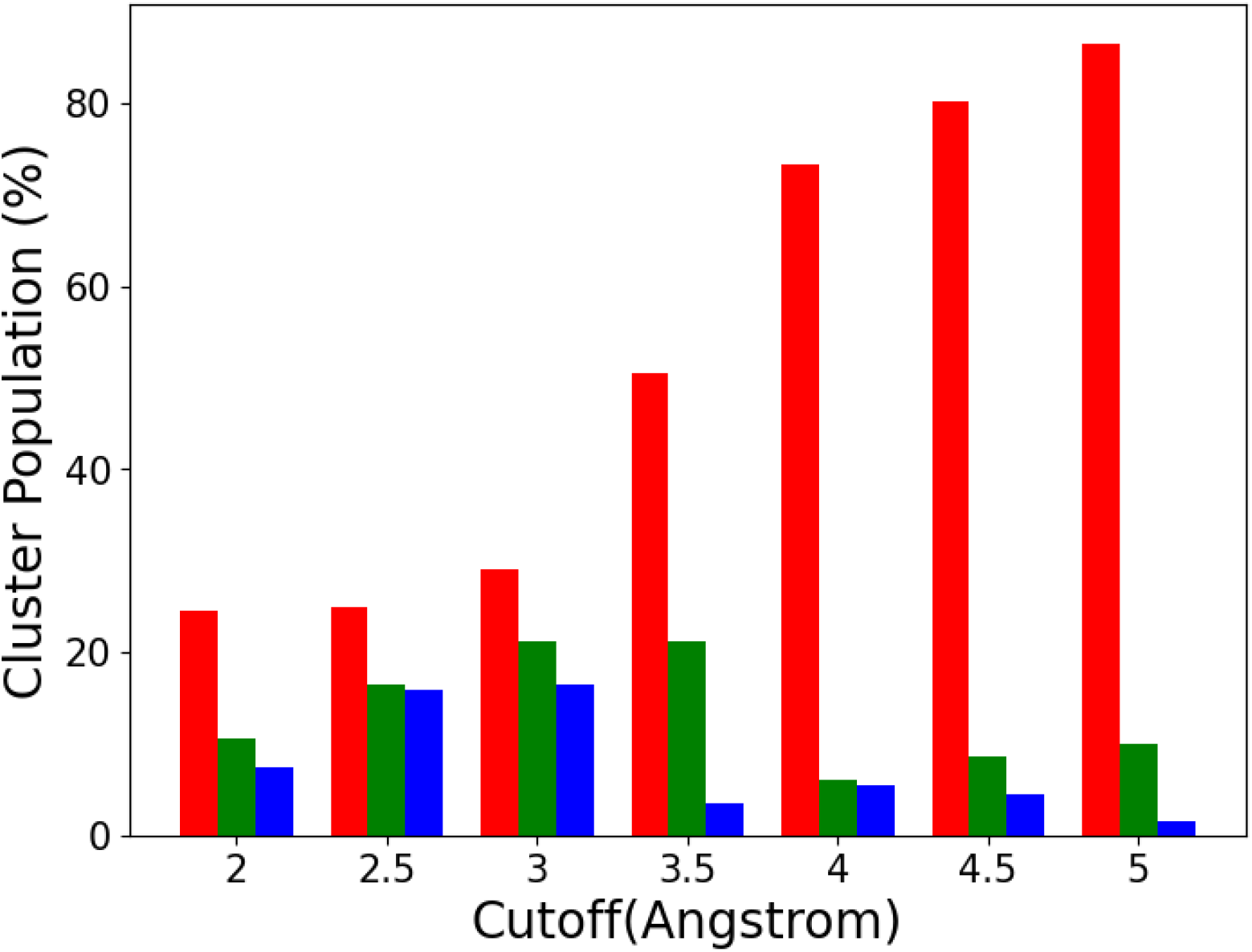
Percentage distribution of the top three most populated structural clusters of the folded state as a function of cutoff distance (Å). The red, green, and blue bars represent the most, second, and third most populated clusters, respectively. As the clustering algorithm uses a similarity basis,^6^ the larger the cutoff, the larger the most populating cluster is, i.e., clustering results are sensitive to the cutoff choice. To minimize this sensitivity, we performed this cluster size analysis by systematically increasing the cutoff and chose the cutoff of 3Å, where the populations of the three most populating clusters are in a plateau.

**Figure S2:**
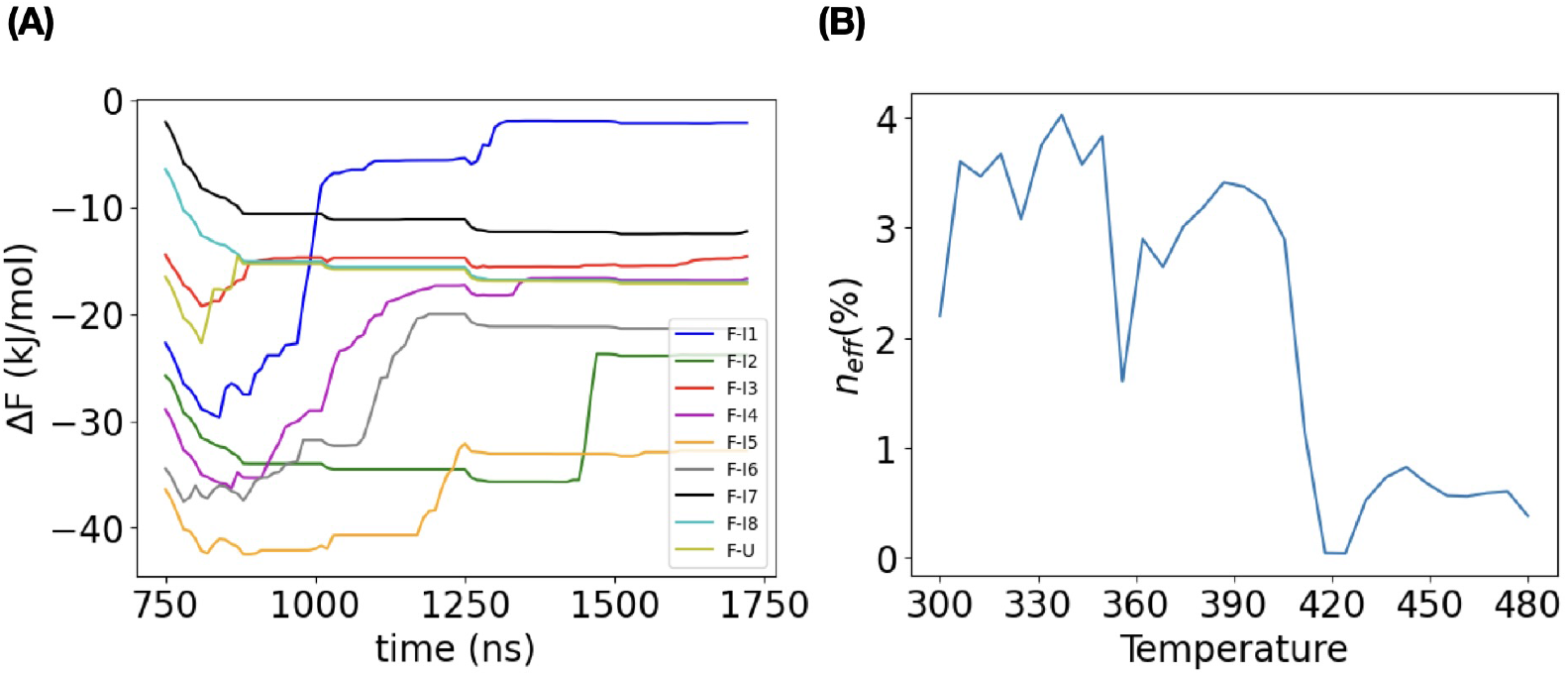
Statistical evaluation for convergence and effective sample size (A) Evolution of free energy differences between any intermediate and unfolded states and the folded state as a function of simulation time. (B) Evaluation of effective sample size (ESS) as a percentage of the total samples.

**Figure S3:**
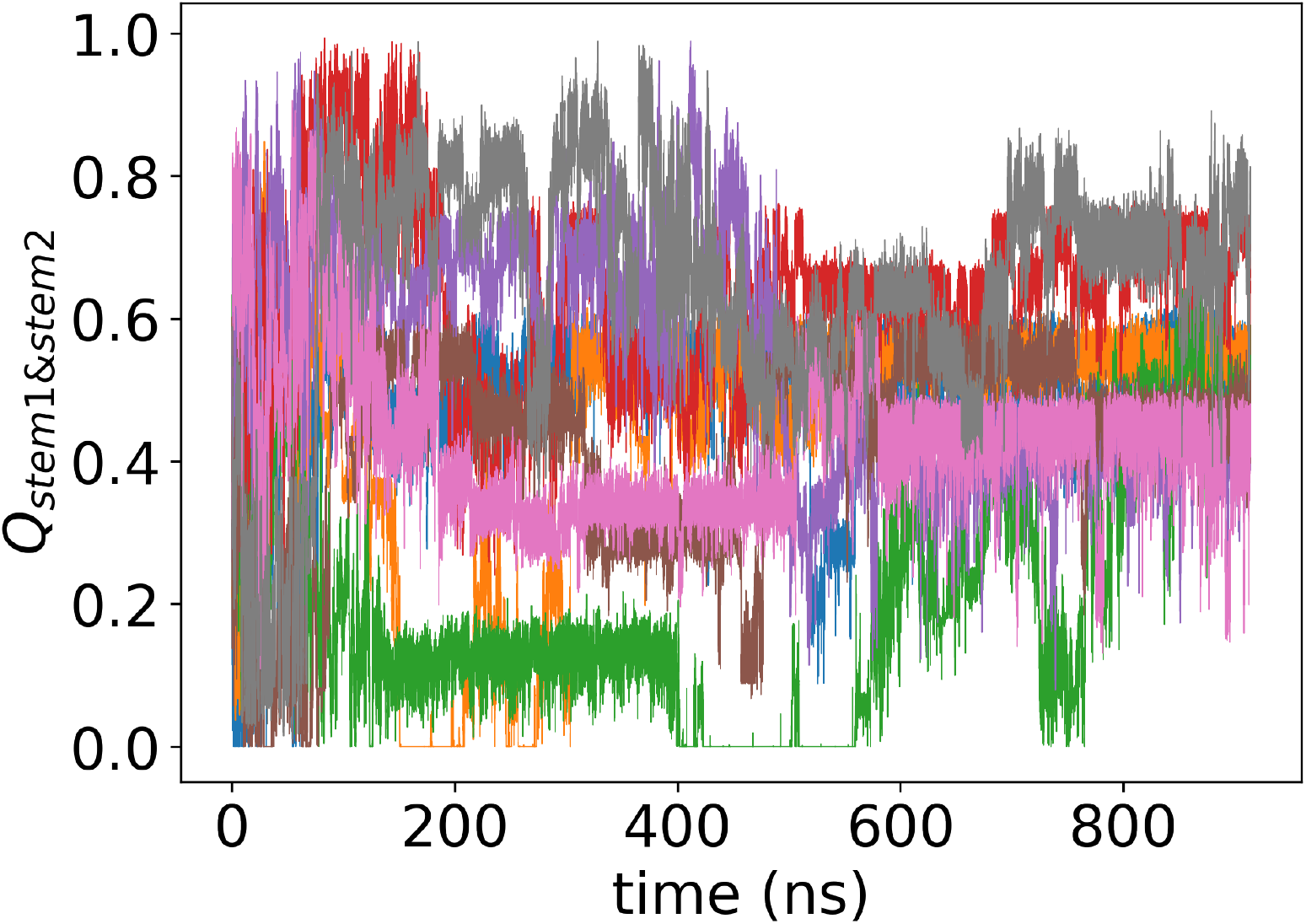
Sampling of *Q*_*stem*1&*stem*2_ as a function of time for all of the walkers (each color represents a different walker). The large regions of white space in the upper and lower portion of the plot indicate that several walkers remained trapped in metastable states for extended periods. In this simulation, the number of native hydrogen bonds in stem 1 and stem 2 (denoted as *N*_*HB*1_, and *N*_*HB*2_) as independent CVs used for biasing and simulation conducted at a temperature range of 300–480*K*, and *T*_*run*_ = 390*K*.

**Figure S4:**
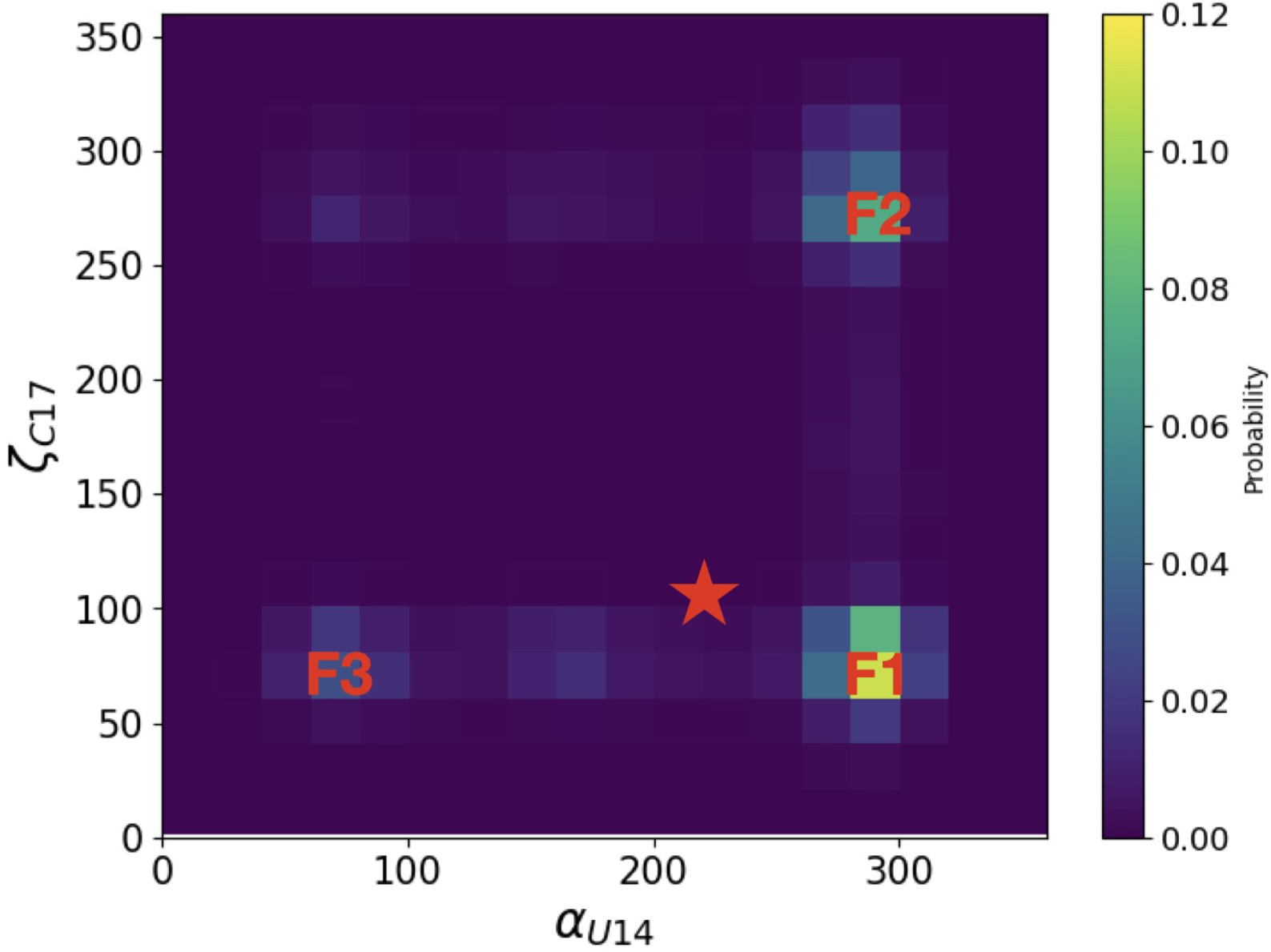
This heatmap illustrates the probability distribution of dihedral angles *α*_*U*14_ (x-axis) and *ζ*_*C*17_ (y-axis) in the folded state of PK1 focusing. The nucleotides U14 and C17 mark the ends of loop 2. The color intensity represents the frequency of conformational occurrences. The red star denotes the native structure (PDB ID: 2G1W), while F1, F2, and F3 locations indicate the angles of the three most populated clusters in the folded state, with F1 being the most populated, followed by F2 and F3. The backbone dihedral angles *α*_*U*14_ and *ζ*_*C*17_ quantify the backbone twisting differences between F1, F2, and F3 best (compared to all other dihedral angles of PK1).

